# Survey of the Phytophthora species diversity reveals *P. abietivora* as a potential Phytophthora root rot pathogen in Québec Christmas tree plantations

**DOI:** 10.1101/2022.10.26.513888

**Authors:** Guillaume Charron, Julie Yergeau, Hervé Van der Heyden, Guillaume J. Bilodeau, Carole Beaulieu, Philippe Tanguay

**Affiliations:** Canada Natural Resources, Laurentian Forestry Centre, 1055 rue Du P.E.P.S., P.O. Box 10380, Québec (Québec), Canada, G1V 4C7; Phytodata Inc. research company, 291 rue de la Coopérative, Sherrington (Québec), Canada J0L 2N0; Département de Biologie, 2500, boulevard de l'Université, Université de Sherbrooke, Sherbrooke (Québec), Canada, J1K 2R1; Canadian Food Inspection Agency, 3851 Fallowfield Road, Ottawa, ON, Canada, K2J 4S1

**Keywords:** Christmas tree disease, ITS1-5.8S-ITS2, Phytophthora isolation, Undescribed *Phytophthora*, Baiting, Oomycetes.

## Abstract

Christmas trees are an economically and culturally important ornamental plant in North America. Many microorganisms are pathogens of firs cultivated as Christmas trees. Among those, *Phytophthora* causes millions of dollars in damage to plantations annually. In Canada, it is still not known which species are responsible for Phytophthora root rot (PRR) of cultivated *Abies* species. Between 2019 and 2021, soil and root samples were collected from 40 Christmas tree plantations in Québec province. We used soil baiting and direct root isolation to assess the diversity of culturable *Phytophthora* spp. The obtained isolates were identified with a multi-locus sequencing approach, and we used the sequencing data to place them along the *Phytophthora* phylogeny. A total of 44 isolates from six different *Phytophthora* species were identified, one fitting the provisional species *P.* sp.□kelmania□. A seventh taxa, represented by a group of 10 isolates, could not be assigned to any known *Phytophthora* species. Among the known species, *Phytophthora abietivora* was the most prevalent isolated species associated with PRR. Pathogenicity trials confirmed the pathogenicity potential of *P. abietivora* on both Fraser and balsam fir seedlings. Our studies provide a first snapshot of the Phytophthora diversity in Québec’s Christmas tree productions and describe multiple potential first associations between *Phytophthora* species and *Abies balsamea* and *A. fraseri*.

## Introduction

With a little less than 1000 km^2^ in cultivation, Christmas trees are an economically important ornamental plant for Canada and the USA (Statistics Canada 2021; USDA National Agricultural Statistics Service 2021). In eastern North America, Fraser fir is the preferred species cultivated as Christmas trees. Reasons for this include shape and growth, combined with its postharvest qualities such as needle retention and durability (Bates et al. 2004; Mitcham-Butler et al. 1988). Unfortunately, Fraser fir is also one of the most susceptible species to Phytophthora root rot (PRR) (Benson and Grand 2000; Chastagner and Benson 2000). Hence, PRR is responsible for millions of dollars (USD) in yearly losses (Grand and Lapp 1974; Kuhlman and Hendrix 1963; Pettersson et al. 2017). The first symptom of PRR is a reddish-brown discoloration of the cambium region of the roots, which is difficult to detect unless roots are examined. As the infection progresses through the root system, cankers may develop on the lower stem, and symptoms such as chlorosis of foliage, wilting of new growth, branch flagging and stunting begin to show in the above-ground parts of the tree (Chastagner and Benson 2000). Unfortunately, it is usually too late to take action at the onset of visible symptoms and infected trees will die within months.

In the last 40 years, most data concerning *Phytophthora* species associated with Christmas tree production came from the USA (Benson and Grand 2000; Chastagner et al. 1990; Grand and Lapp 1974; McKeever and Chastagner 2016). More than 20 *Phytophthora* species have been associated with more than 20 different species of true firs (Farr 2022), with at least 12 *Phytophthora* species known to induce PRR (McKeever and Chastagner 2016). Since the mid-2000s, data available for European countries overlap with what has been found in the USA (Pettersson et al. 2019; Shafizadeh and Kavanagh 2005; Talgø et al. 2006; Talgø et al. 2007; Talgø et al. 2017), suggesting that either the association between these *Phytophthora* species and firs have been longstanding or that, as for other *Phytophthora* species, worldwide plant trade has contributed to their global dissemination (Bienapfl and Balci 2014; Brasier 2013; Jung et al. 2016). While the information currently available connects *Phytophthora* species to PRR symptomatic trees, few studies have shown their pathogenic potential with Koch’s postulates (Frampton and Benson 2012; Hamm and Hansen 1982a; Hoover and Bates 2013; McKeever and Chastagner 2016). When the testing pathogenicity of *Phytophthora*, approaches rely on two main experimental protocols: (i) inoculations, where the pathogen is grown and then applied on the wounded tissues of the host (Petterson 2019, Quesada-Ocampo 2009, Li et al 2019), and (ii) infestation, where the pathogen is added to the potting medium and the infection by zoospores is usually promoted with flooding periods or increased irrigation (Pettersson 2019, McKeever/Chastagner 2016 and 2019, Quesada-Ocampo 2009,). These approaches cover usual infection patterns for soilborne *Phytophthora* species (Hardham 2001; Judelson and Blanco 2005).

The province of Québec represents more than 35% of the total Canadian cultivated area used for Christmas tree plantations. Eighty-five percent of this area can be found in two regions; Estrie and Chaudière-Appalaches (Statistics Canada 2016). The two most cultivated species used as Christmas trees in Québec’s province are Fraser fir (*A. fraseri*) and balsam fir (*A. balsamea*). While Canadian producers had past PRR occurrences in their plantations, the data available on Phytophthora diversity is insufficient to tackle tasks such as developing integrated management practices that could limit the spreading of diseases in these hosts.

The objectives of this study were to explore the diversity of culturable *Phytophthora* species associated with PRR symptomatic Christmas trees in the province of Québec and to assess the pathogenicity potential of the most prevalent species. To attain these objectives, we sampled Christmas tree plantations in Estrie and Chaudière-Appalaches and demonstrated the pathogenic potential of *P. abietivora* isolates obtained from sampling and found in association with both PRR symptomatic *A. balsamea* and *A. fraseri*. Finally, this study constitutes the first documented report for *P. abietivora* in Canada.

## Material and Methods

### Christmas tree plantations sampling

Soils were sampled during the fall of 2019, 2020, and the spring of 2021 from 40 different sites in Chaudière-Appalaches and Estrie (20 sites each, Supplementary Table S1). All the sites were harboring trees showing Phytophthora root rot symptoms including: (i) shoot dieback, (ii) needle reddening, (iii) resinosis, (iv) Persistent dead needles or (v) generalized discoloration of the tree (pale green/ yellowish). Most of the trees presenting symptoms were in zones identified as poorly drained. Symptoms had been present for two to four years in the plantations prior to sampling. In 2019, over the 14 days period leading to the sampling (September 24^th^ to October 7^th^), the region of Estrie registered 50 to 80 mm of accumulated precipitations with maximum temperatures of 15 to 21°C (Agriculture and Agri-Food Canada 2017b, c). The Chaudière-Appalache region registered 50 to 65 mm of accumulated precipitations with maximum temperature of 15 to 18°C during the same period (Agriculture and Agri-Food Canada 2017b, c). In contrast, over the same period leading to the sampling of Chaudière-Appalache in 2020 (September 15^th^ to the 28^th^), 0 to 10 mm of accumulated precipitations were recorded with maximum temperatures of 18 to 27°C (Agriculture and Agri-Food Canada 2017b, c). The Estrie sampling planned in September 2020 was delayed until May 2021 due to COVID-19 travel limitations. In the 14 days period leading to the sampling dates (May 18^th^ to 31^st^), 15 to 35 mm of accumulated precipitation were registered with maximum temperatures between 27 and 30°C (Agriculture and Agri-Food Canada 2017b, c).

**Table 1:**
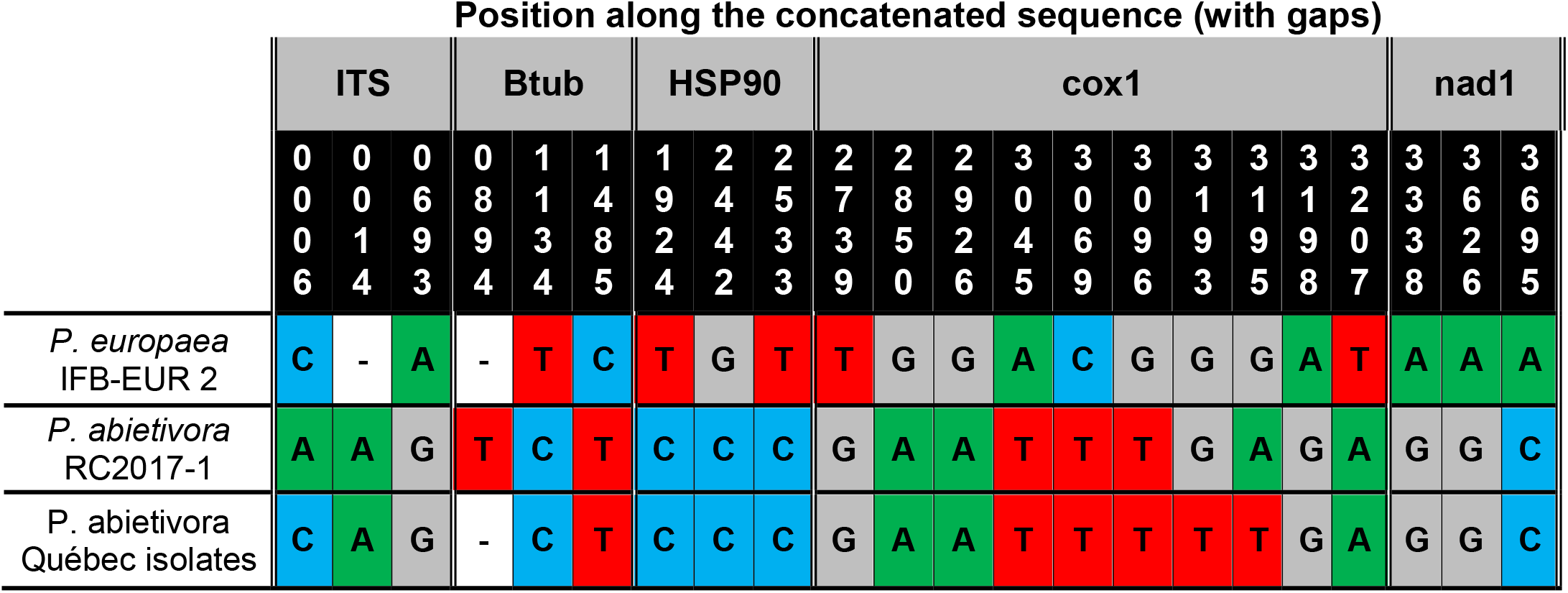
Single nucleotide polymorphisms between *P. abietivora* isolated from Québec and the closest species references

Twelve soil samples were taken in each site from trees aged 4 to 12 years: four showing signs of PRR and four healthy in appearance, and four from apparently healthy trees from a natural forest near the production site. Using a 5-cm diameter soil probe, three soil sub-samples, 120° apart, were taken at the crown limit of the trees up to 17 cm in depth. After removing the organic horizon, the three soil cores were cut in half lengthwise and the three halves were pooled together in a plastic bag (38×19 cm Whirl-Pak) while the remaining halves were put back into the holes left by sampling. Soils samples were identified in numerical order beginning with 001 up to 488. The probe was rinsed with water and heat-disinfected with a propane hand torch between samples. Samples were kept at 4°C until being processed for a minimum of 24 to 72 hours.

### Processing and baiting of soil samples

Each soil sample was mixed and sieved through a 4-mm mesh basket. Root fragments retained in the sieve were transferred at 4°C for oomycetes isolation. Two hundred and fifty milliliters of homogenized soil were put into a clean 1-liter plastic container (IPL plastics, Canada) to which 500 ml of distilled water was added. Four to five healthy rhododendron ’Cunningham’s White’ leaves without any visible injury were rinsed with sterile distilled water and laid upon the water surface of each soil container. Five rhododendron leaves were also placed in a container filled with 500 ml of distilled water to serve as a control. The containers were incubated in the dark at 20°C until the development of hydrosis (water-soaked lesions) on the rhododendron leaf tissue (4 to 7 days).

Following the emergence of hydrosis, leaves were washed with running tap water, dried for 10 minutes on absorbing paper, and taken under a laminar flow hood. Tissues were sampled using a paper punch sterilized with ethanol and flame to obtain leaf disks (6 mm in diameter). Five disks identified A to E were taken at the margin of hydrosis lesions for each soil sample and were inserted into solid V8-PARPNH (clarified V8 50 ml/liter, agar 15 g/liter, PCNB 25 µg/ml, Ampicillin 250 µg/ml, Rifampicin 10 µg/ml, Pimaricin 10 μg/ml, Nystatin 25 µg/ml, and Hymexazol 25 µg/ml) medium (100 mm × 15 mm Petri dishes). Leaf disks were inserted at a 45° angle keeping half the disc above the agar. Petri dishes were incubated in the dark at 20°C and were examined daily for growth. After 2 to 10 days, morphologically distinct colonies were purified by transferring hyphal tips to fresh V8-PARPNH. The cultures were incubated in the dark at 20°C. All cultures not resembling a *Phytophthora* colony were excluded from further investigation. Grown cultures were preserved at 12°C. Isolates were systematically named using a code based on the soil sample they were baited from and the leaf disk they were isolated with. For example: an oomycete obtained from the baiting of soil sample 062 on disk E would be named 062_E. If more than one morphotype was found growing from a leaf disk, a number was appended to the letter (for example: 062_E1 and 062_E2).

### Isolation of oomycetes from roots

Unidentified root fragments gathered from soil samples were transferred to a fine mesh strainer that was first immersed in a beaker containing distilled water and agitated to remove most of the soil. Next, the strainer was then transferred to a beaker containing a 0.5% sodium hypochlorite solution and soaked for 30 seconds, followed by two rinses in distilled water. Surface-sterilized root segments were dried into a laminar flow cabinet for 5 to 10 minutes and cut into 5 to 10 mm sections with a sterile scalpel. Five root sections named A-E were inserted halfway into a V8-PARPNH agar plate. The remaining roots were put into a 2- ml safe-lock tube (Eppendorf, Hamburg, Germany) and stored at −20°C. Media were incubated in the dark at 20°C. Morphotype colonies were purified as described above. Isolate’s name followed the same code used to name soil baiting isolates but an R was added before the soil number (for example: an isolate obtained from a root coming from sample 062 and root section E would be named R_062_E).

### DNA extractions

Purified cultures were grown on V8-agar (1.5% agar, clarified V8 50 ml/liter) overlayed with a sterile cellophane membrane. Mycelia were scraped with a scalpel blade and transferred to 2-ml safelock tubes. A tungsten carbide bead (QIAGEN, Hilden, Germany) was added to each tube and the bottom of the tubes was immersed in liquid nitrogen for 1 minute. The tubes were then quickly transferred to a Mixer mill adapter (Retsch, Haan, Germany) previously cooled at −80°C. Tubes were shaken for 1 minute at a frequency of 26 Hz after which a second cycle of freezing and shaking was performed. DNA was extracted following the protocol developed in Penouilh-Suzette et al. (2020) with minor modifications. In short, 800µl of Lysis buffer (0.1 M Tris-HCl pH 8, 10 mM EDTA pH 8, 1 M NaCl and 1% SDS) were added to up to 50 mg of ground mycelium along with 2 µl of Proteinase K (20 mg/ml). Samples were mixed by inversion and incubated at 50℃ for 10 minutes, while mixing by inversion twice during the incubation. Next, 270 µl of potassium acetate 5 M was added, and tubes were mixed by inversion before incubating on ice for 10 minutes. Samples were centrifuged for 10 minutes (5,000 g), the supernatant was transferred to a new 2 ml tube, and 2 µl of RNaseA (100 mg/ml) was added. The samples were incubated for 30 minutes at 37℃ and mixed by inversion every 10 minutes. Next, 1 ml of Isopropanol was added and tubes were mixed by inversion. The samples were incubated at RT for 30 minutes. Following incubation, samples were centrifuged for 2 minutes at 10,000 × g, the supernatant was removed, and the DNA pellets were rinsed with fresh 70% ethanol. Samples were centrifuged for 1 minute (10,000 × g) and the supernatant was removed. Pellets were air-dried for 1 to 5 minutes under a lamp and then resuspended with 50 µl of PCR-grade water.

### Molecular identification

To identify the isolated organisms, we amplified five loci: the internal transcribed spacers ITS1 and ITS2 including the 5.8S ribosomal RNA region of the rRNA cistron which is referred to as the ITS, the Beta tubulin (Btub), the Heat Shock Protein 90 (HSP90), and the mitochondrial genes Cytochrome oxidase 1 (cox1) and NADH dehydrogenase (nad1). We used primer pairs already described in the literature (Supplementary Table S2). The PCR cycles were: 94□ 2 min; 35 cycles of 94□ 30s; Primer specific Tm 30s; 72□ with an amplicon-specific time; 72□ 10 min. First, PCR products of the expected length for the ITS (Supplementary Table S2) were sent for Sanger sequencing at the CHUL sequencing platform on an ABI 3730xl Data Analyzer (Applied Biosystems, Waltham, USA). Then, for every strain confirmed as *Phytophthora*, the other four loci were amplified and sent for sequencing. For each isolated strain, the sequences were used as a query against the nt/nr collection of the NCBI using BLAST (Altschul et al. 1990). All of the sequences generated for this study were deposited into GenBank under accession numbers ON008003 to ON008178 (for cox1, nad1, Beta-tubulin and HSP90), OM984685 to OM984728 (ITS) and OP676073 (cox1 sequence for the RC2017-1 *P. abietivora* isolate).

### Phylogenetic analysis

For each of the sequenced loci, when two or more isolates had identical sequences, a single sequence was selected to represent the group in the phylogenetic analysis. Sequences of the five loci from strains of 68 *Phytophthora* species belonging to the different known clades were gathered from GenBank and used as references (Supplementary Table S3). Sequences obtained from our isolates and GenBank-downloaded sequences of the reference strains were aligned with Clustal X v2.1 (Larkin et al. 2007) and trimmed using the MEGA X v10.2.6 software (Kumar et al. 2018). For each isolate or reference, the aligned sequences were concatenated using a custom script in R v4.1.1 with the *seqinr* v1.0-2 package (Charif and Lobry 2007; R Core Team 2013) resulting in 3983 sites (with gaps). The phylogenetic analysis employed both maximum-likelihood and Bayesian inference. The maximum-likelihood analysis used the Tamura-Nei model (Tamura and Nei 1993) in MEGA X (Kumar et al. 2018). The dataset comprised 79 concatenated sequences, 68 from the reference strains of the *Phytophthora* species and 11 distinct concatenated sequences groups representing the unique sequences found among the 44 isolates of *Phytophthora* recovered in this study. Positions with less than 90% site coverage were eliminated resulting in 3359 positions in the final dataset. *Elongisporangium undulatum* was designated as an outgroup. Node confidence was estimated with 1000 bootstrap replicates. Bayesian inference was also performed on the dataset using MrBayes3.2.7 (Ronquist et al. 2012). Four Markov chains were used in each of 2 independent runs beginning from random starting trees. The analysis was let run for 13,000,000 generations with tree sampling every 1,000 generations. The first million generations were discarded as burn-in. Out of the remaining trees (24,002), the majority rule consensus tree was computed and branches with 0.95 or more Bayesian posterior probability were considered as significantly supported. The representation of the phylogenetic tree generated by Bayesian inference was represented with Treegraph2 v2.15.0-887 and MEGA X (Kumar et al. 2018; Stöver and Müller 2010).

### Preparation of inoculum and seedlings

Three *Phytophthora abietivora* isolates (062_E, 097_A and 170_B) obtained from the sampling were selected to test their pathogenicity on both balsam and Fraser firs. Isolates were inoculated on a V8- agar medium and incubated for one week at 20°C. Ten mycelial plugs (5 mm ⍰ 5 mm) were taken at the margin of the actively growing cultures and inoculated into a sterile mixture of vermiculite, overnight water-soaked steel-cut oat, and clarified V8 vegetable juice in a 1:1:1 ratio. Controls were prepared the same way using plugs from an uninoculated V8-agar medium. These inocula were incubated for two weeks at 20°C in the dark.

Dormant two-year-old *A. balsamea* and *A. fraseri* container-grown seedlings were obtained from a local nursery (Productions Resinex Inc., La Durantaye, Canada) and kept dormant at −2°C for two months. They were prepared for the experiment by transferring them at 4°C for a week, followed by a week at 10°C. Fraser and balsam fir seedlings were respectively re-potted in individual 1.5-liters and 1-liter plastic containers with pierced bottoms filled with an autoclaved soilless mix containing Pro-mix LP15 peat moss (Premier Tech, Rivière-du-Loup, Canada), vermiculite (Perlite Canada Inc., Lachine, Canada) and silica sand (Atlantic silica inc., Poodiac, Canada) in a 2:1:1 ratio amended with 4 kg/m^3^ of 18-6-8 NPK Nutricote™ fertilizer (Chisso-Asahi Fertilizer Co., Tokyo, Japan). Seedlings were maintained in a greenhouse with a 16-hour photoperiod under high-pressure sodium lighting (≈150 µmol m^-2^s^-1^ at the top of the seedlings) (OSRAM Sylvania Inc., Wilmington, USA), with temperatures of 20°C during the day and 18°C during the night. Seedlings were irrigated for 10 minutes, four times weekly (≈400 ml per irrigation period). These conditions were maintained until bud burst.

### Pathogenicity trials

After bud burst, 100 ml of the top layer of the potting medium was replaced with a fresh potting mix infested with 0.1% (V/V) of the *P. abietivora* inoculum. Trials followed a randomized block design where three blocks containing five replicates of each inoculum/tree species combinations were placed in different areas of the greenhouse (three blocks × two species × (three isolates + one control) × five replicates = 120 seedlings). Growth conditions were the same as stated above except that every two weeks, the seedlings were flooded (1 cm of water above the potting medium) for 48 hours to create conditions favorable to zoospores production and dissemination. The height (soil surface to the tip of the seedling rounded to the closest 5 mm) and the collar diameter of the seedlings were recorded right before inoculation (time point "0"), every other week through the first month (time points 2 and 4), and every week between week five and twelve (time points 5 to 12). Symptoms of PRR were assessed by visual examination of the seedlings looking for stem and branch wilting or chlorotic and dead needles. An estimated percentage of the seedling tissues affected by these symptoms was recorded for each seedling at each time point. Seedlings were considered dead once symptoms affected 100% of their tissues.

After 12 weeks, all seedlings were carefully removed from their containers and the root systems were sampled by separating them from the stem at the height of the root collar. The root systems were washed to remove potting media, retaining as much root tissue as possible. The clean root systems were successively plunged into a beaker containing clean distilled water for 1 minute, then surface sterilized into a beaker containing a 0.5% sodium hypochlorite solution for 30 seconds. Finally, the roots were rinsed twice in clean distilled water for 30 seconds. Surface-sterilized root systems were dried for 10 to 15 minutes in a laminar flow cabinet and then 5 to 10 roots were sampled and cut into 5 to 10-mm segments. Two aliquots of 10 root segments were taken for each seedling. The remainder of the surface sterilized root systems were oven-dried for 48 hours at 70°C. The dried root systems were weighed to estimate the severity of root degradation. We used one of the two aliquots of root segments for oomycete isolation as described in the “Isolation of oomycete from roots” section. The other aliquot was used for DNA extraction using the DNeasy Powersoil Pro Kit (QIAGEN, Hilden, Germany) and molecular detection of *Phytophthora* using the nad1 PCR as described in the “Molecular identification” section.

### Statistical analyses

All statistical analyses were performed with the R software v4.1.1 (R Core Team 2013) using custom scripts and the FSA package v0.9.3 for the Dunn’s Test function (Ogle 2017). Unless stated otherwise, the significance level used for all statistical tests is α = 0.05. The area under the disease progression curves (AUDPC) was calculated as described (Madden et al. 2007). Data representations, survival analyses and statistical comparisons were made using the beanplot v1.3.1 (Kampstra 2008), beeswarm v0.4.0 (Eklund 2021), survival v3.2-11 (Therneau and Grambsch 2000) and survminer v0.4.9 (Kassambra 2021) packages.

## Results

### *Phytophthora* isolations

In 2019, 41 *Phytophthora* isolates were recovered from sampling, while the 2020 sampling yielded three isolates (Supplementary Table S4). Among them, 40 were recovered by baiting soil samples with rhododendron leaves. These isolates were obtained from 22 unique soil samples with some samples yielding up to four isolates. DNA sequencing of the Beta-tubulin, ITS, HSP90, cox1 and nad1 genetic markers coupled with BLAST analyses revealed that the recovered isolates belonged to at least seven species. Based on the ITS sequences, we obtained eight *P. chlamydospora* isolates, eight *P. abietivora* isolates (one from root isolation), seven *P. gonapodyides* isolates, three *P. gregata* isolates, six *P. megasperma* isolates (two from root isolation), and two *P.* sp. □kelmania□ isolates. A group of 10 isolates represented by the isolate 160_A1 (one from root isolation) had discordant BLAST results as their ITS sequence was closest to *P. chlamydospora* while their other four loci sequences did not match *P. chlamydospora*. These isolates were identified as “Undescribed *Phytophthora*” in the context of this study. Except for six isolates of *P. gonapodyides* and two undescribed *Phytophthora* obtained from soil samples taken in natural fir stands in the vicinity of Christmas tree plantations, all other isolates were from soil or root samples gathered under firs showing PRR symptoms. Even though 160 soil samples from healthy trees in plantations were analyzed, no *Phytophthora* was recovered from these samples. The fact that few isolates were obtained from healthy trees in natural forests coupled with the low numbers of *Phytophthora* isolates recovered from 2019, 2020 and 2021 samplings precluded statistical analysis. However, we observed that *P. abietivora* isolates were recovered from the highest number of independent samples (eight isolates from eight independent samples), which could imply a role for this organism in the development of PRR.

### Phylogenetic analysis

Our phylogenetic analysis revealed that the isolates obtained from Christmas tree plantations were related to clade 6, 7 and 8 species (Figure 1). Most of the isolates obtained from our survey were related to known *Phytophthora* species, except for ten 10 undescribed *Phytophthora* isolates (from 7 independent soil samples) that seem related among themselves but not to any other species used as reference. For those sequences, BLAST analyses would not return consistent identity results. Therefore, our phylogeny placed those isolates in clade 6b as a monophyletic taxa sister to the *P. chlamydospora* and *P. gonapodyides* group.

**Fig. 1.**
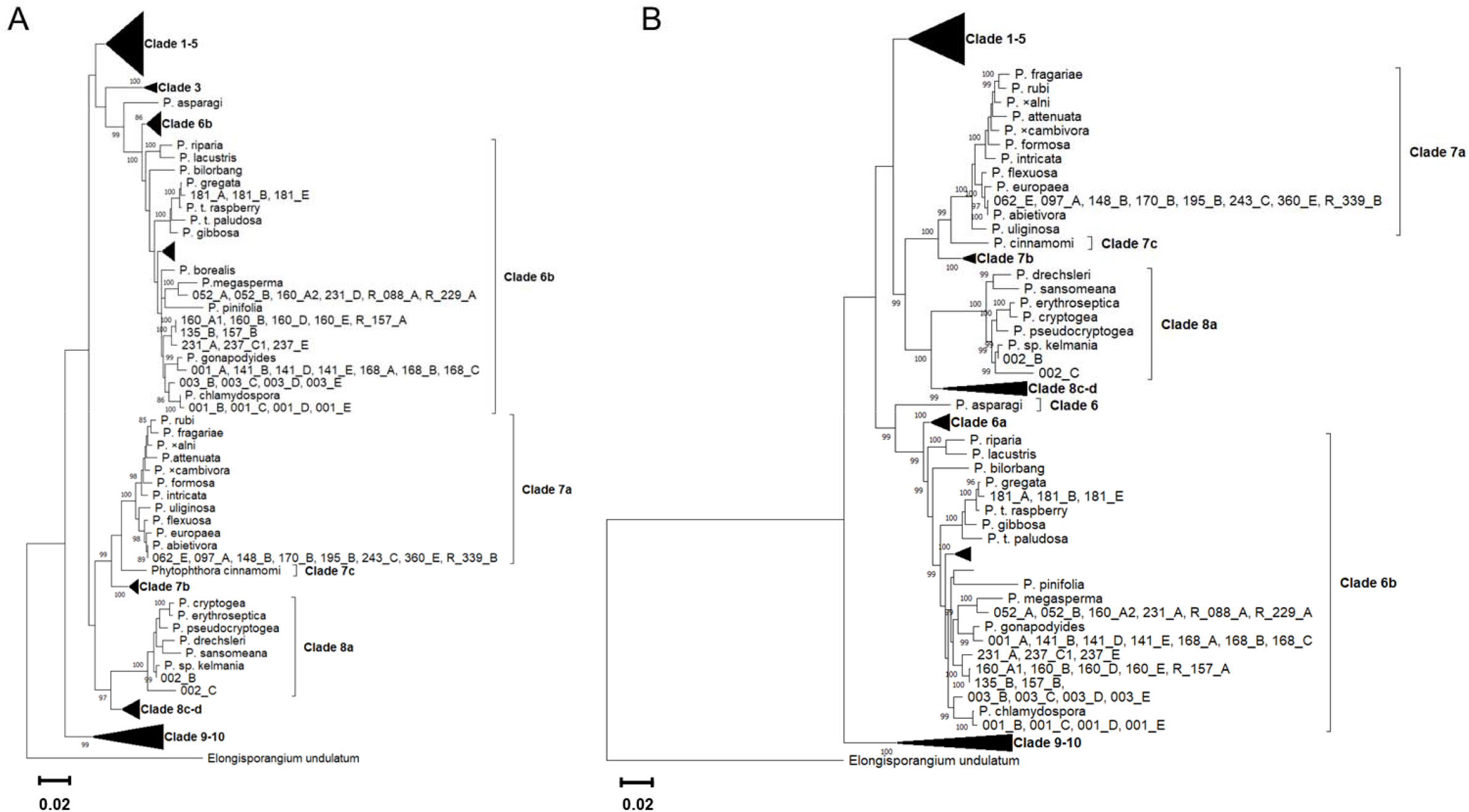
Multiple *Phytophthora* species were recovered in plantations and natural forests: (**A**) Maximum likelihood analysis of the recovered isolates and reference taxa based on concatenated ITS, Cox1, β-Tubulin, Nad1, and HSP90 sequences. *Elongisporangium undulatum* was used as an outgroup. The bootstrap test was conducted with 1,000 replicates bootstrap values >80% are indicated on the nodes. The scale bar shows the number of expected substitutions per site. References are represented by species names and isolates by their isolate code number. (**B**) Bayesian inference analysis of the same sequence alignment. Bayesian posterior probabilities (>95%) are indicated on the nodes.

The sequences from our *P. abietivora* isolates were consistent with the sequences of the *P. abietivora* type strain RC2017-1 with only 3 SNPs (99.93% identity) found over the concatenated sequence data (3973 bp). We identified 22 SNPs (99.45% identity) between our *P. europaea* reference and the *P. abietivora* type strain while there were 20 SNPs (99.50% identity) between our *P. abietivora* isolates and the *P. europaea* reference. Most of the SNPs found were in the cox1 locus (Table 1).

### Progression of PRR and survival of seedlings

Visual examination of the seedlings allowed the observation of a rapid onset of PRR symptoms after the second flooding period (four weeks post-infestation, Figure 2a, b). The statistical comparison of AUDPC values indicated a difference between treatments (Kruskal-Wallis test, H(7) = 118.53, *P* < 0.001). Post-hoc tests showed that, in Fraser fir seedlings, the AUDPC values of infested seedlings significantly differed from the noninfested controls for all the tested isolates (Two-sided Dunn’s test *P* < 0.001, Benjamini-Hochberg corrected). In balsam firs, the results showed that two out of three soil-infested seedling groups significantly differed from the noninfested soil controls (Two-sided Dunn’s test *P*: 170_B = 0.049, 62_E < 0.001, and 97_A = 0.301, Benjamini-Hochberg corrected). The comparison between Fraser and balsam infested seedlings also showed a significant difference in AUDPC (Wilcoxon rank sum test, W=595, *P* < 0.001).

**Fig. 2.**
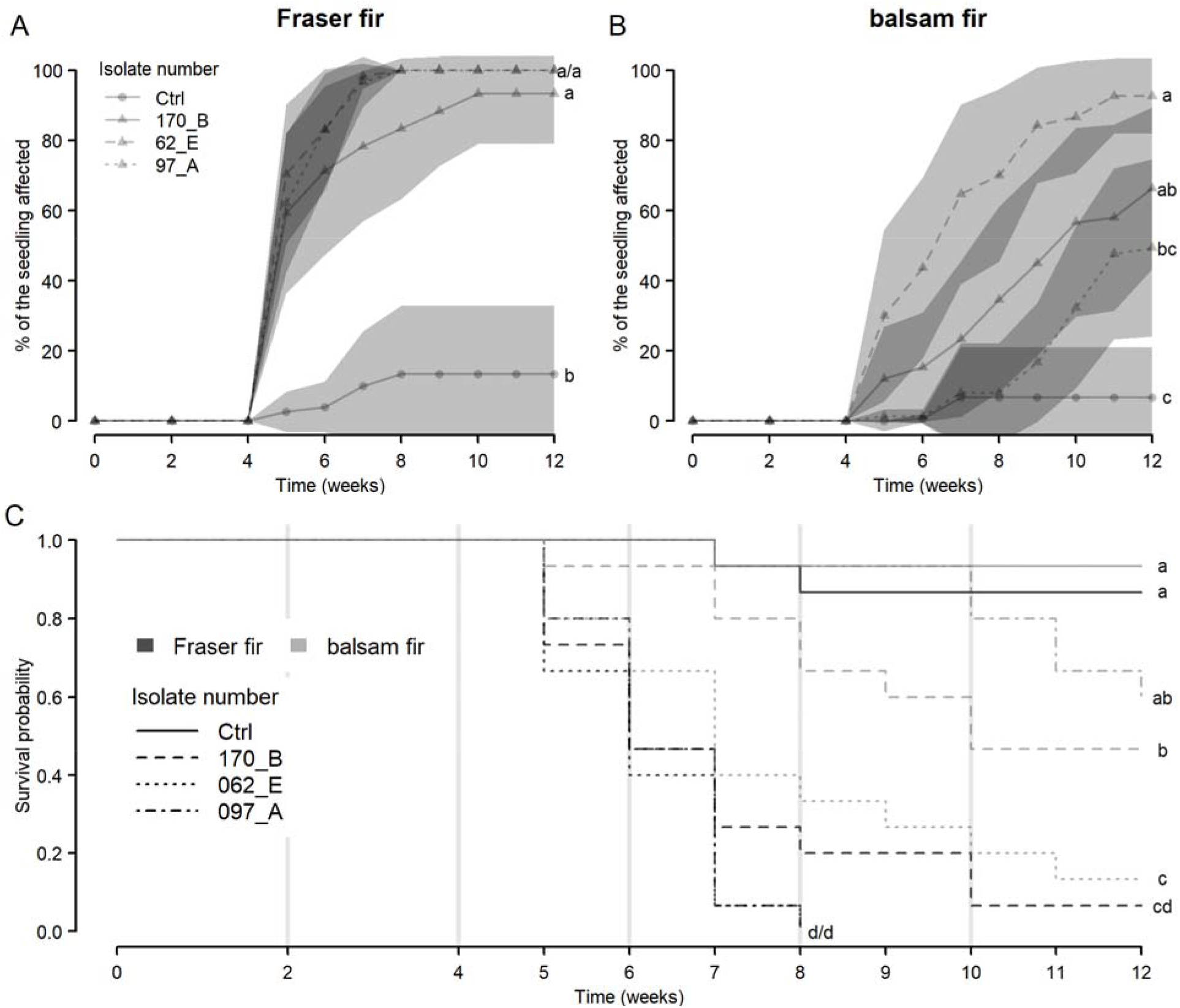
Rapid onset of PRR symptoms lead to seedlings death in inoculated groups: (**A, B**) Disease progression curve in Fraser and balsam fir seedlings represented as the percentage of the plant affected by symptoms of PRR for each seedling. Replicate data was averaged for each tested isolate and non-infested controls (Ctrl); darkened areas represent 95% confidence interval and letters indicate significant differences in AUDPC (two-sided Dunn’s test). (**C**) Survival probability varies between infested and non-infested (Ctrl) seedlings; grey vertical lines represent the timing of each 48-hours flooding period and letters indicate significant differences in survival curves (Log-rank test).

First seedling deaths were recorded as soon as five weeks post-infestation. While noninfested soil controls showed only little losses (Fraser 13.333%, two replicates out of 15 and balsam 6.666%, one replicate out of 15), a large proportion of seedlings in infested soils died (Fraser 97.7%, 44 replicates out of 45 and balsam 60%, 27 replicates out of 45) (Figure 2c). Accordingly, there was a significant difference in survival probability between treatments (log-rank test, X^2^(7) = 95, *P* < 0.001). Pairwise comparisons show that all isolates used to infest soils of Fraser fir seedlings lead to a significantly different survival probability compared to the noninfested soil controls (Figure 2c, log-rank test *P* < 0.001 for all isolates, Benjamini-Hochberg corrected). In balsam fir survival probability was significantly different in two isolates compared to the noninfested soil controls (Figure 2c, log-rank test *P*: 170_B = 0.012, 62_E < 0.001, and 97_A = 0.054 Benjamini-Hochberg corrected). The comparison between Fraser and balsam infested seedling soils showed a significant difference in survival probability. (log-rank test, X^2^(1) = 37.3, *P* < 0.001).

### Impacts of *P. abietivora* isolates on fir seedlings growth

We used the height, stem diameter and root system mass collected during the experiment to evaluate the effect of soil infestation with *P. abietivora* isolates on the growth of balsam and Fraser firs seedlings.

We calculated height increment as the difference in height between the final (t_12_) and initial (t_0_) time points measurements. Statistical comparisons indicated significant differences in height growth between treatments (Kruskal-Wallis test, H(7) = 67.182, *P* < 0.001). Post-hoc tests showed that, in infested Fraser fir soil treatments, seedlings had height increment significantly different than the noninfested soil controls (Figure 3a, Two-sided Dunn’s test *P*: 170_B = 0.040, 62_E = 0.013, and 97_A = 0.004, Benjamini-Hochberg corrected). For balsam fir seedlings, such observations could only be made in soil infested with isolate 62_E (Figure 3b, Two-sided Dunn’s test *P*: 170_B = 0.310, 62_E = 0.04, and 97_A = 0.989, Benjamini-Hochberg corrected).

**Fig. 3.**
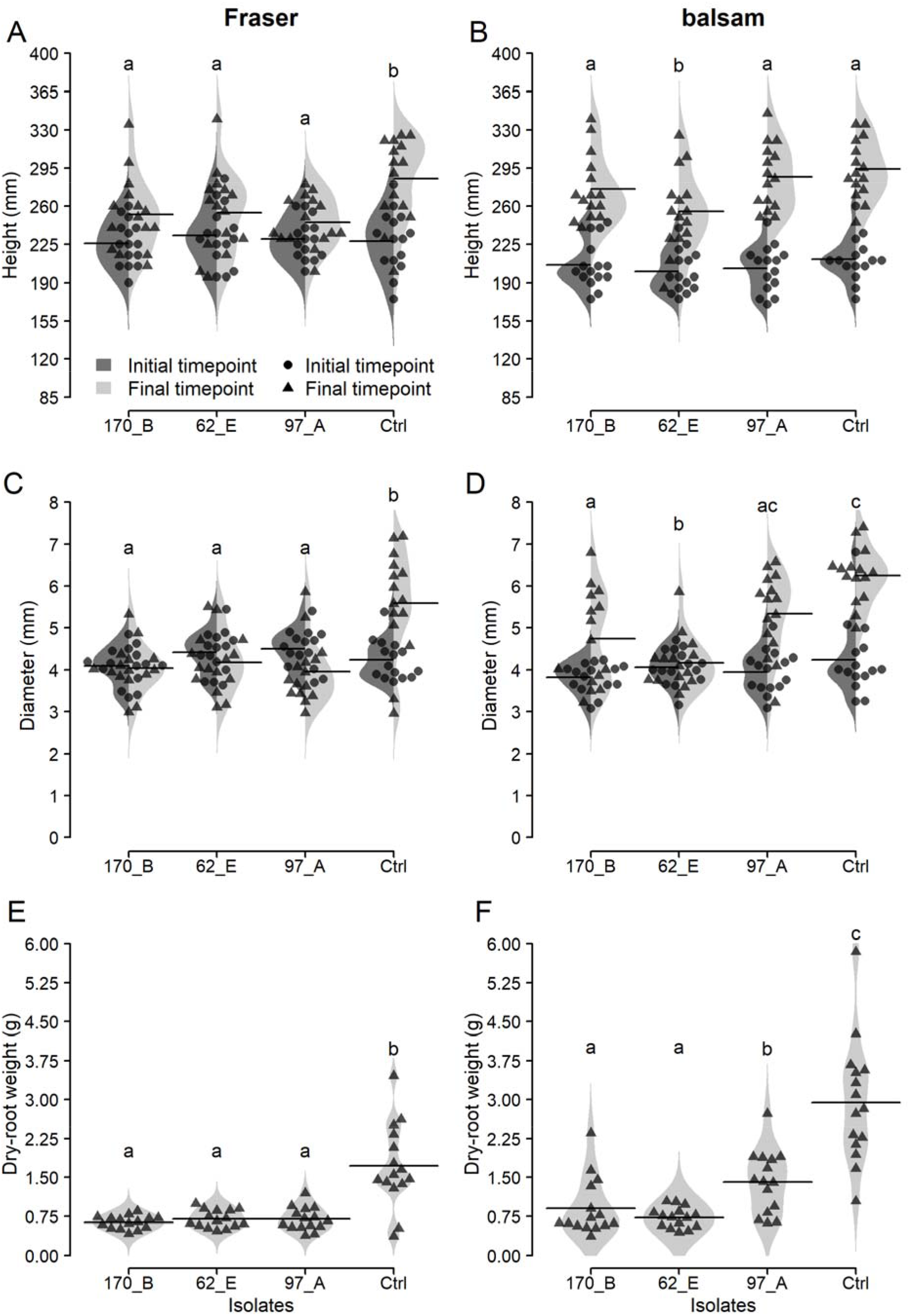
Infestation with *P. europaea*-like isolates affected growth in both Fraser and balsam fir seedlings: (**A, B**) Beanplots comparing height distributions between the initial time point (infestation week, t_0_) and the final time point (12 weeks post-infestation, t_12_) for each isolate inoculated and the non-infested controls (Ctrl). Letters indicate significant differences in height increments (Dunn’s test). (**C, D**) Beanplots comparing collar diameter distribution between t_0_ and t_12_ for each isolate used for infestation and the non-infested controls (Ctrl). Letters indicate significant differences in diameter increments (Dunn’s test). (**E, F**) Beanplots of dry root mass distribution at t_12_ for each isolate used for infestation and the non-infested controls (Ctrl). Letters indicate significant differences in dry root weight (Dunn’s test). Black bars on the plots illustrate the mean of the data. Symbols represent individual data points for each time point.

The root collar diameter increment of the seedling was calculated the same way as the height increment by subtracting the final and initial time points measurements. Statistical comparisons indicated significant differences in root collar diameter increment between treatments (Kruskal-Wallis test, H(7) = 60.946, *P* < 0.001). Post-hoc tests showed that, in soil-infested treatments, Fraser fir seedlings had significantly lower diameter increment than the noninfested soil controls (Figure 3c, Two-sided Dunn’s test *P*: 170_B = 0.003, 62_E < 0.001, and 97_A < 0.001, Benjamini-Hochberg corrected). In balsam fir, only the seedlings infested soils with the 170_B and 62_E *P. abietivora* isolates had significantly different seedling root collar diameter increments than the noninfested soil controls. (Figure 3d, Two-sided Dunn’s test *P*: 170_B = 0.0233, 62_E < 0.001, and 97_A = 0.3189, Benjamini-Hochberg corrected).

Dry-root weight at the final time point also showed significant differences between treatments Kruskal-Wallis test, H(7) = 61.244, *P* < 0.001). Post-hoc tests showed that both soil infested Fraser fir seedlings (Figure 3e, Two-sided Dunn’s test *P*: 170_B < 0.001, 62_E < 0.001, and 97_A = 0.002, Benjamini-Hochberg corrected) and soil infested balsam fir seedlings (Figure 3f, Two-sided Dunn’s test *P*: 170_B = 0.003, 62_E < 0.001, and 97_A = 0.004, Benjamini-Hochberg corrected) had significantly lower dry-root weight than their respective noninfested soil controls.

### Re-isolation of the *P. europaea*-like isolates on diseased seedlings

None of the necrotic root sections used for reisolation of the *P. abietivora* isolates yielded live oomycetes after inoculation on a fresh medium. Using a PCR approach based on the amplification and sequencing of the nad1 locus allowed the detection of the *P. abietivora* isolates in DNA extracted from root samples. While the isolates could be detected in more than a third of our symptomatic or dead seedlings (16/45 for Fraser fir and 18/45 for balsam fir), they were not detected in any of our noninfested soil controls regardless of their health status. Statistical analyses indicate a significant relationship between the positivity of the nad1 PCR and both the status (healthy or symptomatic/dead) of the seedling at the end of the experiment (Pearson’s Chi-squared test with Yates’ continuity correction, X^2^(1, 120) = 16.860, *P* = < 0.001) and its infestation status (Pearson’s Chi-squared test with Yates’ continuity correction, X^2^(1, 120) = 14.008, *P* < 0.001).

## Discussion

Phytophthora root rot has been reported throughout the Christmas fir cultivation range. The problem has been well documented in the USA where both the diversity of the *Phytophthora* species (Chastagner and Benson 2000; McKeever and Chastagner 2016; Wisconsin Department of Agriculture 2015) and the severity (Benson and Grand 2000) of the PRR were described. Surveys were also conducted in European countries to identify *Phytophthora* species associated with PRR on Christmas trees (Pettersson et al. 2019; Shafizadeh and Kavanagh 2005; Talgø et al. 2006; Talgø et al. 2007). While PRR is present in Canadian Christmas tree plantations, details about the cohorts *Phytophthora* species and the intensity of the disease are lacking in Canada. To fulfill this gap, we surveyed PRR in Christmas tree plantations in Quebec, the province with the largest production in Canada.

It is important to take into consideration that baiting, like other enrichment culture methods, will introduce biases in the diversity of observed isolated species depending on many factors such as baiting method, bait used, isolation medium used, and moment chosen for baiting (Burgess et al. 2017; Davison and Tay 2005; Erwin and Ribeiro 1996). This will often lead to an underestimation of diversity. Therefore, our present study is by no means a comprehensive report on the Phytophthora diversity found in the Christmas tree plantations, but rather a first exploration step. Approaches such as metabarcoding are usually complementary to baiting studies as they allow a more complete picture of the diversity present in the field instead of only the culturable one (La Spada et al. 2022). They also offer opportunities for better comparisons between the different environments sampled using quantitative measures such as diversity indices. To that end we extracted the DNA from all the soil, baited leaves, and root samples used to isolate Phytophthora, allowing future metabarcoding analyses.

Despite extensive surveying over three years, only a few *Phytophthora* isolates were recovered. In total, 44 *Phytophthora* isolates belonging to seven different taxa were recovered. Of the 480 soil samples processed, 22 (4.58%) yielded at least one *Phytophthora* isolate. This is low compared to other studies that reported *Phytophthora* recovery rates of 21% from forest soil samples of the Eastern and North-Central U. S. oak ecosystems (Balci et al. 2007), 50% for the Fraser fir plantations in the Southern Appalachians (Pettersson et al. 2017), and 9% from the total sampled trees in North Carolina (Benson and Grand 2000). Our *Phytophthora* spp. recovery rate was still lower when isolation from PRR symptomatic trees was considered. Fifteen soil samples (12.55%) yielded *Phytophthora* isolates compared to 54% and 20% respectively reported by Pettersson et al. (2017) and Benson and Grand (2000). Direct isolation from unidentified root fragments was generally less successful than baiting, with less than 10% (4 out of 44) of the isolates obtained from root samples. However, there was a certain degree of incertitude about the origins of the root fragments as they were sampled from the soil near the trees and not directly cut from the root systems. This could, in part, explain our lower isolation successes. Similar results were obtained by (Aghighi et al. 2016), where most of the *Phytophthora* isolates associated with declining *Rubus anglocandicans* were recovered by baiting rather than direct isolation from roots. As most of our isolates were obtained from the 2019 sampling, questions arise about the sampling success difference between sampling years. Looking at the climatic data available, 2020 and 2021 had very dry growing seasons. At the time of sampling, the departure from average precipitation varied from −55 mm to −170 mm in the sampled areas for 2020 and −15 mm to −80 mm in 2021 (Agriculture and Agri-Food Canada 2017a). Comparatively, the 2019 growing season had a positive departure from average precipitation. Moreover, the Estrie sampling was postponed from fall 2020 to spring 2021 due to COVID-19 restrictions. We think these climatic differences could have decreased the relative abundance of *Phytophthora* species on the sampled sites. This hypothesis could be confirmed using future metabarcoding data.

*Phytophthora chlamydospora*, P. *gonapodyides*, and *P. gregata* belong to clade 6, and are considered riparian-associated species (Brasier et al. 1993; Jung et al. 2011). These three species have been reported worldwide in various natural ecosystems (Hansen et al. 2015; Wan et al. 2020). *Phytophthora chlamydospora* and *P. gonapodyides*, are believed to mainly act as saprophytes, with episodic fine root infections on weakened hosts (Hansen et al. 2015). Associations between *Phytophthora gonapodyides* and true fir species were previously reported (Chastagner et al. 1995; McKeever and Chastagner 2016; Pettersson et al. 2019), but to our knowledge, this study is the first to report the association between *P. gonapodyides* and balsam fir (*A. balsamea*). *Phytophthora chlamydospora* was previously found to be associated with *A. concolor* and *A. procera* (Brasier et al. 1993). Still, the present study is the first to report associations of this species with *A. balsamea* and *A. fraseri*. *Phytophthora gregata* was identified as an aggressive pathogen of Australian wetland plants, being able to infect through the roots and the stem of susceptible hosts (Wan et al. 2020). Other studies have reported associations of *P. gregata* with other broadleaf forest species (see the summary table of global *Phytophthora gregata* records in Wan et al. 2020). Still, to our knowledge, this study is the first report of this species in Canada, and the first to report the association of this species with a conifer species (*A. balsamea*). Further studies are required to investigate the pathogenicity of *P. gregata* on true firs.

Unexpectedly, we observed a group of 10 undescribed *Phytophthora* isolates with ITS sequence close to *P. chlamydospora* but had discordant results for the other loci. These isolates may represent a new species that could be confirmed with more in-depth genomic and taxonomic studies.

*Phytophthora megasperma* represents a broad, unresolved species complex (Hansen et al. 1986; Hansen et al. 2009) with some members having broad host ranges (Hansen and Maxwell 1991). Isolates identified as *P. megasperma* are known pathogens of Douglas-fir (*Pseudotsuga menziesii*) (Hamm and Hansen 1982b). *P. megasperma* isolates have also been isolated from *A. fraseri*, *A. grandis*, and *A. procera* in the USA (Chastagner et al. 1995; McKeever and Chastagner 2016), from *A. procera* in Ireland (Shafizadeh and Kavanagh 2005), from *A. lasiocarpa* in Norway (Talgø et al. 2007), and *A. nordmanniana* in Sweden (Pettersson et al. 2019). This study is the first to report associations of *P. megasperma* with *A. balsamea*.

True firs were shown to be susceptible to *Phytophthora* sp. □kelmania□, but this study is the first to report the association of this *Phytophthora* species with balsam fir (*A. balsamea*). Previously, its DNA was detected in Wisconsin from diseased Canaan, Fraser and Douglas fir root samples (Wisconsin Department of Agriculture 2015). McKeever and Chastagner (2016) have isolated *P.* sp. □kelmania□ from Fraser fir (*A. fraseri*), Canaan fir (*A. balsamea var. phanerolepis*) and Turkish fir (*A. bornmuelleriana*). This was the second most isolated *Phytophthora* species in North Carolina on Fraser fir showing PRR symptoms, and artificial inoculations confirmed that both balsam and Fraser fir were highly susceptible to *P.* sp. □kelmania□. McKeever and Chastagner (2019) have shown that the aggressiveness of P. sp. *kelmania* and other *Phytophthora* species on Fraser and balsam firs is partially temperature dependent. More recently, Kondo et al. (2022) performed artificial inoculations and showed that *P.* sp. □kelmania□ induced PRR symptoms in Sakhalin fir seedlings (*A. sachalinensis*). While there is limited literature available about the isolation of this species, GenBank lists sequences for more than 40 isolates (including the two presented in this study). Even with such a small sample size, more than half of these isolates have been isolated from firs in eastern North America. When Molnar et al. (2020) revisited the historical strain collection of the Pennsylvania Department of Agriculture, *P*. sp. □kelmania□ was one of the most commonly isolated species, and it was associated with counties with big Christmas tree plantations (Molnar et al. 2020). Moreover, consistent *P*. sp. □kelmania□ recovery could be traced back as early as 1988. Based on these observations, *P*. sp. □kelmania□ seems to be a fir pathogen endemic to the east coast. Formally naming and describing this species should be undertaken as soon as possible to benefit researchers and Christmas tree producers.

Our phylogenetic analysis revealed that our isolates were closely related to *P. abietivora*. However, our phylogeny and that of Li et al. (2019), which formally described *P. abietivora*, show some discordances. They show *P. abietivora* as a sister species to *P. flexuosa* and those two as a sister group to *P. europaea* (Li et al. 2019) while we show *P. europaea* and *P. abietivora* in a sister group to *P. flexuosa*. This may result from a difference in the resolution of both phylogenies as ours covers most of the *Phytophthora* clades but lacks multiple sequences per species, and Li’s phylogeny focuses on clade 7 species. While *P. abietivora* is a very recently described species initially supported by a single isolate (Li et al. 2019), the high sequence homology (99.93% identity) to the type strain leaves little doubt on the identity of our isolates. This study represents the first report of *P. abietivora* in Canada and its presence in soils associated with *A. balsamea* showing PRR-like above-ground symptoms in Canadian Christmas tree plantations.

*Phytophthora europaea* was initially isolated from soil associated with declining oak forests (*Quercus spp.* and *Q. robur)* in France and Germany (Jung et al. 2002). It has since been isolated in Europe and associated with *Quercus* species (Balci and Halmschlager 2003; Tkaczyk et al. 2017), in North America associated with diverse deciduous tree species and riparian ecosystems (Balci et al. 2006; Reed et al. 2019; Reeser et al. 2011; Sims et al. 2015), and in Australia associated with *Dacrydium cupressinum* and *Iris sibirica* (International Collection Of Microorganisms (ICMP) 2022). More recently, surveys of Christmas tree plantations in the USA recovered anecdotal *P. europaea* isolates on *A. fraseri* (McKeever and Chastagner 2016; Pettersson et al. 2017). *P. europaea* has been reported twice in Canada, once from a diseased balsam fir imported from North Carolina (Robideau et al. 2011), and a second time from a diseased balsam fir reported in the Annual 2017 Canadian Plant Disease Survey (Elmhirst 2017).

Given their high sequence similarity (≈99.5% identity), isolates were probably identified as *P. europaea* in studies preceding the formal description of *P. abietivora* and would now need to be reconsidered as *P. abietivora*. Molnar et al. (2020) reported the identification of *P. abietivora* isolates from the collection of the Pennsylvania Department of Agriculture. They could track the first isolation of *P.abietivora* from *A. fraseri* and *Tsuga canadensis* as far back as 1989. They also discuss the high similarity in the sequences with *P. europaea*, especially for the ITS (Molnar et al. 2020). More recently, Bily et al. (2022) isolated *P. abietivora* from multiple deciduous trees, which would fit with the *P. europaea* hosts already described. They also observed genetic variation between their strains and the *P. abietivora* type strain for the loci they sequenced (Bily et al. 2022). In California, a survey reported the isolation of *Phytophthora* sp. *cadmea,* baited from the soil of restoration areas. This undescribed taxon has high sequence similarity with *P. europaea* and *P. abietivora* (Bourret et al. 2020). Based on those observations, we suggest caution when considering *P. europaea* and *P. abietivora* as different species. While we agree that these species may represent diverging lineages, more extensive population genomics and phenotypic studies may be necessary to untangle the relationship between them. It is interesting to note that to date *P. europaea* is not reported as associated with *Abies* species in Europe. Moreover, it is considered a weak pathogen of deciduous trees (Jung et al. 2017). In North America, Bily et al. (2022) reported that *P. abietivora* strains recently collected on deciduous trees also seemed to be weak pathogens to the deciduous trees species they challenged with their strains. Considering this information, an interesting hypothesis could be that *P. europaea* was introduced to North America from Europe through plant nursery trade like other species from this genus (Goss et al. 2009; Moralejo et al. 2009). After spreading throughout North America, it could have invaded the Christmas tree plantation environment and started diverging, becoming the *P. abietivora* that was recently described. Population genomic studies of *P. europaea /P. abietivora* strains from Europe and North America could help us confirm this scenario.

Pathogenicity trials conducted with three *P. abietivora* isolates obtained from soil associated with diseased trees confirmed that this species has the potential to infect both *A. balsamea* and *A. fraseri* seedlings. However, disease progression and survival curves suggest that the infection in *A. balsamea* progresses slower than in *A. fraseri*, suggesting that the *P. abietivora* isolates are more aggressive on *A. fraseri*. The sensitivity of *A. fraseri* to PRR is well documented for other *Phytophthora* species (Benson and Grand 2000; Hinesley et al. 2000; Quesada-Ocampo et al. 2009) and could explain this difference in survival throughout our greenhouse experiment. However, as McKeever and Chastagner (2019) showed that aggressiveness of *Phytophthora* species on firs is usually lower at the temperatures used in this study; the difference in mortality observed may be an overestimate. It is difficult to explain why our reisolation attempts from root tissues were unsuccessful when the isolates’ DNA could be recovered in root tissues. One hypothesis could be that our surface sterilization was too harsh, but our methodology did not differ much from what is typically used in such experiments (Bily et al. 2022; Pettersson et al. 2017). While we could not reisolate live oomycetes from the roots of dead seedlings, the fact that we only detected the DNA of the isolates used to infest the soil in the roots of seedlings showing symptoms of PRR strongly suggests that the isolates can be involved in the development of the disease observed in our seedlings.

This study provides a sample of the Phytophthora diversity present in the Christmas tree cultures of the province of Québec. For many of the species reported here, the association with either *A. fraseri* or *A. balsamea* is novel information and many were not reported yet in Canada. More investigation will be needed to evaluate the impact of these potential pathogens as our pathogenicity trials were only conducted on the *P. abietivora* isolates and Koch’s first and fourth postulates were not formally verified. We also present strong evidence that *P. abietivora* isolates are potential root rot pathogens of balsam fir and confirm their pathogenicity on Fraser firs. The information we provide will be helpful in the development of targeted detection tools and pathogen management strategies that will help producers lower the risks of PRR outbreaks in their plantations. A follow-up study, involving metabarcoding is also underway to establish a more complete portrait of Phytophthora and oomycetes diversity in the various samples (soil, baiting and root) collected in this study.

## Acknowledgments

The authors would like to thank Jean-François Légaré, Éric Dussault, Christian Lacroix, Dominique Choquette and Jacinthe Drouin for their contributions to fieldwork and sampling; Dr Christopher I. Keeling for comments on the manuscript; Marie-Krystel Gauthier, Marie-Josée Bergeron and Véronique Lévesque-Tremblay for their precious technical advice; Michel Gravel, Marquis Goupil, Renald Gilbert, Claire Gilbert, Larry Downey, Christian Vanasse, Serge Vaillancourt, Yohan Blanchette, Junior Beloin, Gérald Couture, Jacques Fortin and David Gouin for giving us access to their plantations and the Ministère de l’agriculture des pêcheries et de l’alimentation du Québec for providing funding for this study.

