## Supplementary Table 1-4 for "Survey of the Phytophthora species diversity reveals *P. abietivora* as a potential Phytophthora root rot pathogen in Québec Christmas tree plantations"

**Supplementary table S1.** List of surveyed Christmas tree plantations

| Site number | Year(s) | Location | GPS coordinates | Planted fir species | Type of soil | Weed management | Symptomatic tree % | Soil sample ID |
|---|---|---|---|---|---|---|---|---|
| 1 | 2019 and 2020 | Chaudière-Appalaches | 46.714471, -71.212684 | balsam | Sandy loam to loamy sand | unknown | 5-10 | 001-012 and 337-356 |
| 2 | 2019 and 2020 | Chaudière-Appalaches | 46.780222, -71.120361 | balsam | Loamy sand | unknown | 5-15 (2019)<br>1-10 (2020)[a] | 013-024 and 357-368 |
| 3 | 2019 | Chaudière-Appalaches | 46.171528, -70.848972 | balsam | Loam to sandy loam | unknown | 1-5 | 025-036 |
| 4 | 2019 | Chaudière-Appalaches | 46.068222, -70.929111 | balsam "cook" | Loam | unknown | > 25 | 037-048 |
| 5 | 2019 | Chaudière-Appalaches | 46.056972, -71.124556 | balsam "cook" | Loam | unknown | > 25 | 049-060 |
| 6 | 2019 | Chaudière-Appalaches | 45.991139, -71.485611 | balsam | Loam to sandy loam | unknown | 5-10 | 061-072 |
| 7 | 2019 | Chaudière-Appalaches | 45.954488, -70.858245 | balsam | unknown | unknown | 1-10 | 073-084 |
| 8 | 2019 | Chaudière-Appalaches | 45.954492, -70.858245 | balsam "cook" | unknown | unknown | 1-5 | 085-096 |
| 9 | 2019 | Chaudière-Appalaches | 45.938861, -70.856917 | balsam | Loam to silty loam | unknown | 1-5 | 097-108 |
| 10 | 2019 | Chaudière-Appalaches | 45.938861, -70.856917 | balsam | Loam to silty loam | unknown | 1-5 | 109-120 |
| 11 | 2019 and 2021 | Estrie | 45.172889, -71.487605 | balsam | Loam | Good (2019)<br>Poor (2021) | 5 (2019)<br>< 5 (2021)[a] | 181-192 and 369-380 |
| 12 | 2019 and 2021 | Estrie | 45.230506, -71.937717<br>45.231361, -71.927792 | Fraser (2019)<br>balsam (2021) | Loam | Good | 5 (2019)<br>< 5 (2021)[a] | 121-132 and 381-392 |
| 13 | 2019 and 2021 | Estrie | 45.114804, -71.998998 | balsam | Loam | Average | 5 (2019)<br>< 5 (2021)[a] | 133-144 and 453-464 |
| 14 | 2019 | Estrie | 45.036552, -72.021969 | Fraser | Loam | Average | < 5 | 145-156 |
| 15 | 2019 and 2021 | Estrie | 45.063416, -71.875822 | balsam | Loam | Average | 20 (2019)<br>10 (2021)[a] | 157-168 and 429-440 |
| 16 | 2019 and 2021 | Estrie | 45.230306, -71.764462 | Fraser | Sandy loam | Average | 20 (2019)<br>15 (2021)[a] | 169-180 and 393-404 |
| 17 | 2019 and 2021 | Estrie | 45.355868, -71.829743 | balsam | Clay loam | Average | 5 (2019)<br>< 5 (2021)[a] | 193-204 and 417-428 |

| Site number | Year(s) | Location | GPS coordinates | Planted fir species | Type of soil | Weed management | Symptomatic tree % | Soil sample ID |
|---|---|---|---|---|---|---|---|---|
| 18 | 2019 and 2021 | Estrie | 45.373306, -71.697270 | balsam | Sandy loam | Average | 10 (2019)<br>5-10 (2021)[a] | 205-216 and 405-416 |
| 19 | 2019 | Estrie | 45.674133, -71.147980 | balsam | Loam to silty loam | Good | < 5 | 217-228 |
| 20 | 2019 and 2021 | Estrie | 45.902924, -71.023241 | Fraser | Loam to sandy loam | Good | 5 (2019)<br>< 5 (2021)[a] | 229-240 and 465-476 |
| 21 | 2020 | Chaudière-Appalaches | 45.9550, -70.82834 | balsam | Loam to silty loam | Good | 1-5 | 241-252 |
| 22 | 2020 | Chaudière-Appalaches | 46.02861, -70.87861 | Fraser | Loam | Good | 1-5 | 253-264 |
| 23 | 2020 | Chaudière-Appalaches | 46.01888, -70.87527 | Fraser | Loam | Good | 1-5 | 265-276 |
| 24 | 2020 | Chaudière-Appalaches | 45.96583, -70.79666 | balsam | unknown | Average | unknown | 277-288 |
| 25 | 2020 | Chaudière-Appalaches | 45.87638, -70.8572 | balsam | Loam to sandy loam | Good | 1-5 | 289-300 |
| 26 | 2020 | Chaudière-Appalaches | 45.92416, -70.84666 | balsam "cook" | Loam to silty loam | Average | 1-5 | 301-312 |
| 27 | 2020 | Chaudière-Appalaches | 45.93305, -70.75375 | balsam | Loam to silty loam | Good | 1-5 | 313-324 |
| 28 | 2020 | Chaudière-Appalaches | 46.05916, -70.80722 | balsam | Loam | Good | 1-5 | 325-336 |
| 29 | 2021 | Estrie | 45.077082, -72.043458 | Fraser | Loam | Average | < 5 | 441-452 |
| 30 | 2021 | Estrie | 45.761968, -70.993923 | Fraser | Loam | Good | < 5 | 477-488 |

[a]Decrease in prevalence due to the removal of affected trees observed the previous year.

**Supplementary table S2**. Primers detailed information

| Oligo name | Sequence | Target | Tm (℃) | Elongation time | Expected length (bp) | Reference |
|---|---|---|---|---|---|---|
| Btub_F1 | 5'-GCCAAGTTCTGGGAGGTCATC-3' | Beta-Tubulin gene | 54.0 | 1 min 30 s | 1228 | [1] |
| Btub_R1 | 5'-CCTGGTACTGCTGGTACTCAG-3' | | | | | [2] |
| HSP90_F1 | 5'-GCTGGACACGGACAAGAACC-3' | Heat-shock protein 90 | 56.8 | 1 min | 921 | [1] |
| HSP90_R1 | 5'-ACACCCTTGACRAACGACAG-3' | | | | | |
| NADH_F1 | 5'-CTGTGGCTTATTTTACTTTAG-3' | NADH dehydrogenase subunit 1 | 51.1 | 1 min | 897 | [2] |
| NADH_R1 | 5'-CAGCAGTATACAAAAACCAAC-3' | | | | | |
| ITS5_F | 5'-GGAAGTAAAAGTCGTAACAAGG-3' | Internal transcribed spacer 1 - 5.8S - Internal transcribed spacer 2 | 50.0 | 1 min | 944 | [3] |
| ITS4_R | 5'-TCCTCCGCTTATTGATATGC-3' | | | | | |
| Fm85mod | 5'-TCAWCWMGATGGCTTTTTTCAAC-3' | Cytochrome c oxidase subunit I | 54.0[a] | 1 min | 774 | [4] |
| OOmCoxILevup | 5'-RRHWACKTGACTDATRATACCAAA-3' | | | | | |

[a] 1-minute annealing time

[1] Blair, J. E., Coffey, M. D., Park, S.-Y., Geiser, D. M., and Kang, S. 2008. A multi-locus phylogeny for Phytophthora utilizing markers derived from complete genome sequences. Fungal Genetics and Biology 45:266-277.

[2] Kroon, L. P. N. M., Bakker, F. T., van den Bosch, G. B. M., Bonants, P. J. M., and Flier, W. G. 2004. Phylogenetic analysis of Phytophthora species based on mitochondrial and nuclear DNA sequences. Fungal Genetics and Biology 41:766-782.

[3] White, T. J., Bruns, T., Lee, S., and Taylor, J. 1990. Amplification and direct sequencing of fungal ribosomal RNA genes for phylogenetics. PCR protocols: a guide to methods and applications 18:315-322.

[4] Robideau, G. P., De Cock, A. W., Coffey, M. D., Voglmayr, H., Brouwer, H., Bala, K., Chitty, D. W., Désaulniers, N., Eggertson, Q. A., Gachon, C. M., Hu, C. H., Küpper, F. C., Rintoul, T. L., Sarhan, E., Verstappen, E. C., Zhang, Y., Bonants, P. J., Ristaino, J. B., and Lévesque, C. A. 2011. DNA barcoding of oomycetes with cytochrome c oxidase subunit I and internal transcribed spacer. Mol Ecol Resour 11:1002-1011

**Supplementary table S3.** References sequences used for phylogenetic analyses

| Species | Clade | Genbank Accession # | | | | |
|---|---|---|---|---|---|---|
| | | ITS | NADH | CoxI | HSP90 | Btub |
| *P. idaei* | 1a | AF266773.1 | AY564012.1 | GU945479.1 | EU080133.1 | AY564070.1 |
| *P. tentaculata* | 1b | AJ854302.1 | AY564031.1 | AB688300.1 | EU080157.1 | AY564089.1 |
| *P. infestans* | 1c | MG865514.1 | AY563979.1 | MH136908.1 | MK020322.1 | MH493956.1 |
| *P. phaseoli* | 1c | AF266778.1 | AY563986.1 | AY129168.1 | EU079917.1 | AY564044.1 |
| *P. botryosa* | 2a | DQ275188.1 | AY563993.1 | HQ261257.1 | EU079938.1 | AY564051.1 |
| *P. capsici* | 2b | MG865467.1 | DQ361203.1 | AB688176.1 | MK020283.1 | MH493915.1 |
| *P. citricola* | 2c | AY670705.1 | AY563997.1 | FJ237512.1 | MK020288.1 | AY564055.1 |
| *P. frigida* | 2d | KX011269.1 | KX011293.1 | MN991993.1 | KX011241.1 | MN991988.1 |
| *P. ilicis* | 3 | AF339443.1 | AY564013.1 | AY129172.1 | EU079863.1 | EU079860.1 |
| *P. nemorosa* | 3 | KF317082.1 | DQ361211.1 | HQ261374.1 | KX250968.1 | KX250965.1 |
| *P. quercina* | 3b | FJ801390.1 | DQ361213.1 | HQ261406.1 | EU080493.1 | EU080490.1 |
| *P. megakarya* | 4 | AF266782.1 | AY564020.1 | HQ261356.1 | EU079973.1 | AY564078.1 |
| *P. agathidicida* | 5 | KP295308.1 | KP295334.1 | KP295222.1 | KP295276.1 | KX251077.1 |
| *P. cocois* | 5 | KP295306.1 | KP295356.1 | KP295244.1 | KP295298.1 | KX251105.1 |
| *P. heveae* | 5 | AF266770.1 | AY564009.1 | HQ261320.1 | EU079734.1 | AY564067.1 |
| *P. asparagi* | 6 | EU301167.2 | JN547679.1 | HQ012844.1 | HQ012890.1 | JN547591.1 |
| *P. gemini* | 6a | NR_147866.1 | MF326932.1 | MH620038.1 | KX251129.1 | KX251126.1 |
| *P. humicola* | 6a | AF266792.2 | AY564011.1 | AY564184.1 | EU080172.1 | AY564069.1 |
| *P. inundata* | 6a | EF210201.1 | JN936043.1 | EF210207.1 | JN935947.1 | EF210203.1 |
| *P. rosacearum* | 6a | HQ012958.1 | JN936032.1 | HQ012882.1 | HQ012926.1 | JN935980.1 |
| *P. t.* walnut | 6a | KC291550.1 | JN936042.1 | JN935971.1 | JN935956.1 | JN935990.1 |
| *P. amnicola* | 6b | JQ029958.1 | JQ029942.1 | JQ029950.1 | JQ029946.1 | JQ029954.1 |
| *P. bilorbang* | 6b | JQ256377.1 | JQ256378.1 | JQ256375.1 | JQ256376.1 | JQ256374.1 |
| *P. fluvialis* | 6b | EU593261.2 | JN547681.1 | JF701440.1 | JF701437.1 | JN547593.1_ |
| *P. gibbosa* | 6b | HQ012933.1 | JN547683.1 | HQ012846.1 | HQ012892.1 | JN547596.1 |
| *P. gonapodyides* | 6b | MG865501.1 | JN936025.1 | MH136896.1 | EU080122.1 | MH493944.1 |
| *P. gregata* | 6b | EU301171.2 | JN547688.1 | HQ012851.1 | HQ012897.1 | JN547600.1 |
| *P. lacustris* | 6b | HQ012956.1 | JN547706.1 | HQ012880.1 | HQ012924.1 | JN547618.1 |
| *P. litoralis* | 6b | EU869199.1 | JN547697.1 | HQ012865.1 | HQ012910.1 | JN547610.1 |
| *P. megasperma* | 6b | EU301166.2 | AY564021.1 | AY564194.1 | HQ012912.1 | AY564079.1 |
| *P. pinifolia* | 6b | EU725806.1 | JN936030.1 | GU799673.1 | N935950.1 | GU799661.1 |
| *P. riparia* | 6b | JQ626595.1 | KF606718.1 | JQ626621.1 | KX251351.1 | JQ626607.1 |
| *P. t.* paludosa | 6b | HQ012953.1 | JN547702.1 | HQ012876.1 | HQ012920.1 | JN547615.1 |
| *P. t.* raspberry | 6b | EU240068.1 | JN936040.1 | JN935969.1 | JN935954.1 | JN935988.1 |
| *P. thermophila* | 6b | HQ012952.1 | JN547699.1 | HQ012871.1 | HQ012915.1 | JN547612.1 |
| *P. borealis* | 6b | MH620128.1 | KF606712.1 | MH136854.1 | KX251191.1 | KX251188.1 |
| *P. chlamydospora* | 6b | MG865471.1 | MN883607.1[a] | MH136867.1 | MK020285.1 | MH493919.1 |
| *P. alni* | 7a | EU048808.1 | DQ202491.1 | GU945463.1 | EU079597.1 | EU079594.1 |
| *P. attenuata* | 7a | KU899196.1 | KU899516.1 | KU899351.1 | KU899431.1 | KU899274.1 |
| *P. cambivora* | 7a | AJ007040.1 | DQ202492.1 | DQ202501.1 | EU080462.1 | EU080460.1 |
| *P. europaea* | 7a | AF449493.1 | AB649211.1 | HQ261303.1 | EU079485.1 | EU079482.1 |
| *P. flexuosa* | 7a | KU517152.1 | KU899544.1 | KU517146.1 | KU899459.1 | KU899302.1 |
| *P. formosa* | 7a | KU517153.1 | KU899512.1 | KU517147.1 | KU899427.1 | KU899270.1 |
| *P. fragariae* | 7a | FJ801768.1 | KU899548.1 | HQ261311.1 | EU079747.1 | EU079744.1 |
| *P. intricata* | 7a | KU899202.1 | KU899523.1 | KU899357.1 | KU899438.1 | KU899281.1 |
| *P. rubi* | 7a | GU259113.1 | FJ696621.1 | HQ261413.1 | KU899391.1 | KU899234.1 |
| *P. uliginosa* | 7a | HQ261721.1 | KU899471.1 | HQ261468.1 | EU080014.1 | EU080012.1 |
| *P. abietivora* | 7a | MK163944.1 | MK164269.1 | OP676073.1[b] | MK164275.1 | MK164274.1 |
| *P. melonis* | 7b | AB366548.1 | EF372622.1 | AY659698.1 | MK020341.1 | AY659687.1 |

| | | | | | | |
|---|---|---|---|---|---|---|
| *P. niederhauserii* | 7b | GU230789.1 | GU477619.1 | GU477617.1 | KU899389.1 | GU477613.1 |
| *P. vignae* | 7b | L41388.1 | AY564032.1 | AY564205.1 | MK087047.1 | AY564090.1 |
| *P. cinnamomi* | 7c | AF087478.1 | AY563996.1 | AY564169.1 | KU899390.1 | AY564054.1 |
| *P. cryptogea* | 8a | MG865483.1 | AY564001.1 | AY564174.1 | MK020297.1 | AY564059.1 |
| *P. drechsleri* | 8a | KJ744314.1 | AY564002.1 | AY564175.1 | MK020299.1 | AY564060.1 |
| *P. erythroseptica* | 8a | AY659445.1 | AY564003.1 | AY659585.1 | MK020302.1 | AY659491.1 |
| *P. sp.kelmania* | 8a | FJ801461.1 | EU497932.1 | HQ261439.1 | EU079609.1 | EU079606.1 |
| *P. pseudocryptogea* | 8a | KP288373.1 | KP288356.1 | KP288339.1 | MK020376.1 | KP288389.1 |
| *P. sansomeana* | 8a | FJ966880.1 | FJ966877.1 | FJ966881.1 | MK020390.1 | FJ966879.1 |
| *P. hibernalis* | 8c | AY423295.1 | AY564010.1 | AY564183.1 | MK020315.1 | AY564068.1 |
| *P. lateralis* | 8c | AY848937.1 | AY564018.1 | AY564191.1 | MK020330.1 | AY564076.1 |
| *P. ramorum* | 8c | MT031967.1 | AY564034.1 | MT235266.1 | MK020385.1 | AY564092.1 |
| *P. syringae* | 8d | AY787032.1 | AY564030.1 | AY564203.1 | MK020393.1 | AY564088.1 |
| *P. hydrogena* | 9a | KC249959.1 | KU695496.1 | MH136902.1 | KX252284.1 | KX252281.1 |
| *P. captiosa* | 9b | FJ801493.1 | MG543021.1 | MG543005.1 | EU079669.1 | EU079666.1 |
| *P. fallax* | 9b | FJ801495.1 | MG543017.1 | GU594811.1 | EU080038.1 | EU080035.1 |
| *P. boehmeriae* | 10 | AB367384.1 | AY563992.1 | HQ261251.1 | EU080165.1 | EU080162.1 |
| *P. richardiae* | 10a | AF271221.1 | AY564028.1 | AY564201.1 | MK020386.1 | AY564086.1 |
| *Elongisporangium undulatum* | outgroup | AY436638.1 | DQ361224.1 | HQ708987.1 | EU080443.1 | DQ361165.1 |

[a] Extracted from the Mitochondrion complete sequence.
[b] Sequenced in this study

**Supplementary table S4.** List of isolated strains

| Isolate | Site number | Associated tree species | Status | Isolation method | Phylogeny closest species | Sequence Accession # | | | | |
|---|---|---|---|---|---|---|---|---|---|---|
| | | | | | | ITS | NADH | CoxI | HSP90 | Btub |
| 001_A | 1 | *A. balsamea* | PRR | Baiting | *P. gonapodyides* | OM984685 | ON008047 | ON008003 | ON008135 | ON008091 |
| 001_B | 1 | *A. balsamea* | PRR | Baiting | *P. chlamydospora* | OM984686 | ON008048 | ON008004 | ON008136 | ON008092 |
| 001_C | 1 | *A. balsamea* | PRR | Baiting | *P. chlamydospora* | OM984687 | ON008049 | ON008005 | ON008137 | ON008093 |
| 001_D | 1 | *A. balsamea* | PRR | Baiting | *P. chlamydospora* | OM984688 | ON008050 | ON008006 | ON008138 | ON008094 |
| 001_E | 1 | *A. balsamea* | PRR | Baiting | *P. chlamydospora* | OM984689 | ON008051 | ON008007 | ON008139 | ON008095 |
| 002_B | 1 | *A. balsamea* | PRR | Baiting | *P.* sp. "kelmania" | OM984690 | ON008052 | ON008008 | ON008140 | ON008096 |
| 002_C | 1 | *A. balsamea* | PRR | Baiting | *P.* sp. "kelmania" | OM984691 | ON008053 | ON008009 | ON008141 | ON008097 |
| 003_B | 1 | *A. balsamea* | PRR | Baiting | *P. chlamydospora* | OM984692 | ON008054 | ON008010 | ON008142 | ON008098 |
| 003_C | 1 | *A. balsamea* | PRR | Baiting | *P. chlamydospora* | OM984693 | ON008055 | ON008011 | ON008143 | ON008099 |
| 003_D | 1 | *A. balsamea* | PRR | Baiting | *P. chlamydospora* | OM984694 | ON008056 | ON008012 | ON008144 | ON008100 |
| 003_E | 1 | *A. balsamea* | PRR | Baiting | *P. chlamydospora* | OM984695 | ON008057 | ON008013 | ON008145 | ON008101 |
| 052_A | 5 | *A. balsamea* 'Cooks' | PRR | Baiting | *P. megasperma* | OM984696 | ON008058 | ON008014 | ON008146 | ON008102 |
| 052_B | 5 | *A. balsamea* 'Cooks' | PRR | Baiting | *P. megasperma* | OM984697 | ON008059 | ON008015 | ON008147 | ON008103 |
| 062_E | 6 | *A. balsamea* | PRR | Baiting | *P. abietivora* | OM984698 | ON008060 | ON008016 | ON008148 | ON008104 |
| 097_A | 9 | *A. balsamea* | PRR | Baiting | *P. abietivora* | OM984699 | ON008061 | ON008017 | ON008149 | ON008105 |
| 135_B | 13 | *A. balsamea* | PRR | Baiting | Undescribed *Phytophthora* | OM984700 | ON008062 | ON008018 | ON008150 | ON008106 |
| 141_B | 13 | *A. balsamea* | Forest | Baiting | *P. gonapodyides* | OM984701 | ON008063 | ON008019 | ON008151 | ON008107 |
| 141_D | 13 | *A. balsamea* | Forest | Baiting | *P. gonapodyides* | OM984702 | ON008064 | ON008020 | ON008152 | ON008108 |
| 141_E | 13 | *A. balsamea* | Forest | Baiting | *P. gonapodyides* | OM984703 | ON008065 | ON008021 | ON008153 | ON008109 |
| 148_B | 14 | *A. fraserii* | PRR | Baiting | *P. abietivora* | OM984704 | ON008066 | ON008022 | ON008154 | ON008110 |
| 157_B | 15 | *A. balsamea* | PRR | Baiting | Undescribed *Phytophthora* | OM984705 | ON008067 | ON008023 | ON008155 | ON008111 |
| 160_A1 | 15 | *A. balsamea* | PRR | Baiting | Undescribed *Phytophthora* | OM984706 | ON008068 | ON008024 | ON008156 | ON008112 |
| 160_A2 | 15 | *A. balsamea* | PRR | Baiting | *P. megasperma* | OM984707 | ON008069 | ON008025 | ON008157 | ON008113 |
| 160_B | 15 | *A. balsamea* | PRR | Baiting | Undescribed *Phytophthora* | OM984708 | ON008070 | ON008026 | ON008158 | ON008114 |

| Isolate | Site number | Associated tree species | Status | Isolation method | Phylogeny closest species | ITS | NADH | CoxI | HSP90 | Btub |
|---|---|---|---|---|---|---|---|---|---|---|
| 160_D | 15 | *A. balsamea* | PRR | Baiting | Undescribed *Phytophthora* | OM984709 | ON008071 | ON008027 | ON008159 | ON008115 |
| 160_E | 15 | *A. balsamea* | PRR | Baiting | Undescribed *Phytophthora* | OM984710 | ON008072 | ON008028 | ON008160 | ON008116 |
| 168_A | 15 | *A. balsamea* | Forest | Baiting | *P. gonapodyides* | OM984711 | ON008073 | ON008029 | ON008161 | ON008117 |
| 168_B | 15 | *A. balsamea* | Forest | Baiting | *P. gonapodyides* | OM984712 | ON008074 | ON008030 | ON008162 | ON008118 |
| 168_C | 15 | *A. balsamea* | Forest | Baiting | *P. gonapodyides* | OM984713 | ON008075 | ON008031 | ON008163 | ON008119 |
| 170_B | 16 | *A. fraserii* | PRR | Baiting | *P. abietivora* | OM984714 | ON008076 | ON008032 | ON008164 | ON008120 |
| 181_A | 11 | *A. balsamea* | PRR | Baiting | *P. gregata* | OM984715 | ON008077 | ON008033 | ON008165 | ON008121 |
| 181_B | 11 | *A. balsamea* | PRR | Baiting | *P. gregata* | OM984716 | ON008078 | ON008034 | ON008166 | ON008122 |
| 181_E | 11 | *A. balsamea* | PRR | Baiting | *P. gregata* | OM984717 | ON008079 | ON008035 | ON008167 | ON008123 |
| 195_B | 17 | *A. balsamea* | PRR | Baiting | *P. abietivora* | OM984718 | ON008080 | ON008036 | ON008168 | ON008124 |
| 231_A | 20 | *A. fraserii* | PRR | Baiting | Undescribed *Phytophthora* | OM984719 | ON008081 | ON008037 | ON008169 | ON008125 |
| 231_D | 20 | *A. fraserii* | PRR | Baiting | *P. megasperma* | OM984720 | ON008082 | ON008038 | ON008170 | ON008126 |
| 237_C1 | 20 | *A. balsamea* | Forest | Baiting | Undescribed *Phytophthora* | OM984721 | ON008083 | ON008039 | ON008171 | ON008127 |
| 237_E | 20 | *A. balsamea* | Forest | Baiting | Undescribed *Phytophthora* | OM984722 | ON008084 | ON008040 | ON008172 | ON008128 |
| R_088_A | 8 | *A. balsamea* 'Cooks' | PRR | Root isolation | *P. megasperma* | OM984725 | ON008087 | ON008043 | ON008175 | ON008129 |
| R_157_A | 15 | *A. balsamea* | PRR | Root isolation | Undescribed *Phytophthora* | OM984726 | ON008088 | ON008044 | ON008176 | ON008130 |
| R_229_A | 20 | *A. fraserii* | PRR | Root isolation | *P. megasperma* | OM984727 | ON008089 | ON008045 | ON008177 | ON008131 |
| 243_C | 21 | *A. balsamea* | PRR | Baiting | *P. abietivora* | OM984723 | ON008085 | ON008041 | ON008173 | ON008132 |
| 360_E | 2 | *A. balsamea* | PRR | Baiting | *P. abietivora* | OM984724 | ON008086 | ON008042 | ON008174 | ON008133 |
| R_339_B | 1 | *A. balsamea* | PRR | Root isolation | *P. abietivora* | OM984728 | ON008090 | ON008046 | ON008178 | ON008134 |
